# A Multiplex Pharmacogenetics Assay using the MinION Nanopore Sequencing Device

**DOI:** 10.1101/563262

**Authors:** Yusmiati Liau, Simone L. Cree, Simran Maggo, Allison L. Miller, John F. Pearson, Patrick A. Gladding, Martin A. Kennedy

## Abstract

**Aim:** The MinION nanopore sequencing device opens the opportunity to cost-effective and point-of-care DNA sequencing. We developed a multiplex assay targeting pharmacogenetic variants related to clopidogrel and warfarin, two commonly used drugs that show response variability due to genetic polymorphisms.

**Materials & Methods:** Six reference and 78 clinical DNA samples were amplified by PCR to generate 15 amplicons targeting key variants. These products were then barcoded to enable sample multiplexing. Three variant calling tools were used to compare genotyping accuracy.

**Results and Conclusions:** All but three samples were successfully sequenced and genotyped. Nanopolish software achieved accuracy > 90 % for all except one variant. While minor mis-genotyping issues exist, this work demonstrates that drug-specific or broad pharmacogenetic screening assays are possible on the MinION sequencing device.

## Introduction

Clopidogrel and warfarin are two commonly used drugs for anticoagulation, with high variability in inter-individual responses. For both drugs, much of the variability is attributed to genetic factors, with pharmacokinetic variation in patients taking clopidogrel being associated with variation in the *CYP2C19* gene [1] and warfarin with *CYP2C9* and *VKORC1* [2]. Both *CYP2C19* and *CYP2C9* are polymorphic genes with more than 30 known alleles [1]. Moreover, recent publications suggest that other genes are also associated with variable drug response, including *B4GALT2* [3] and *ABCB1* [4, 5] for clopidogrel and *GGCX* [6] and *CYP4F2* [6] for warfarin.

Various assays have been developed to screen clopidogrel and warfarin related variants including point of care tests [7, 8], that usually detect a subset of common variants. Extensive panels of pharmacogenetic variants are available through technologies such as mass spectrometry or next generation sequencing. These approaches, however, involve high capital cost and are not always accessible by small-scale laboratories.

MinION (Oxford Nanopore Technologies (ONT), Oxford, UK) is a pocket-sized, portable sequencing device, using the disruptive nanopore technology [9]. It contains an array of nano-scale protein pores embedded in a membrane, across which a current is passed, and single stranded DNA or RNA can be streamed through the pores and sequenced in real-time [10, 11]. The device offers the advantage of portability and almost-zero capital cost, opening the opportunity for point-of-care sequencing, particularly given recent release of smaller-scale, cheaper flow cells (“flongles”, ONT). The main factor limiting its wide utility is the relatively high error rate compared to other sequencing technologies. However, the technology is evolving rapidly, with improvements in the device, chemistries and bioinformatic tools, which are contributing to improvements in accuracy [12].

As proof-of-principle for a potential point of care test, we developed a multiplex assay on this technology, focusing on pharmacogenetic variants related to variability in clopidogrel and warfarin response. The assay contained genetic variants reported as clinically relevant by the Clinical Pharmacogenomics Implementation Consortium (CPIC) guidelines [13], as well as covering new variants reported in recent publications [3-6, 15] (Table 1). Use of a polymerase chain reaction (PCR) barcoding approach enabled us to multiplex 84 samples in one sequencing run.

**Table 1.**
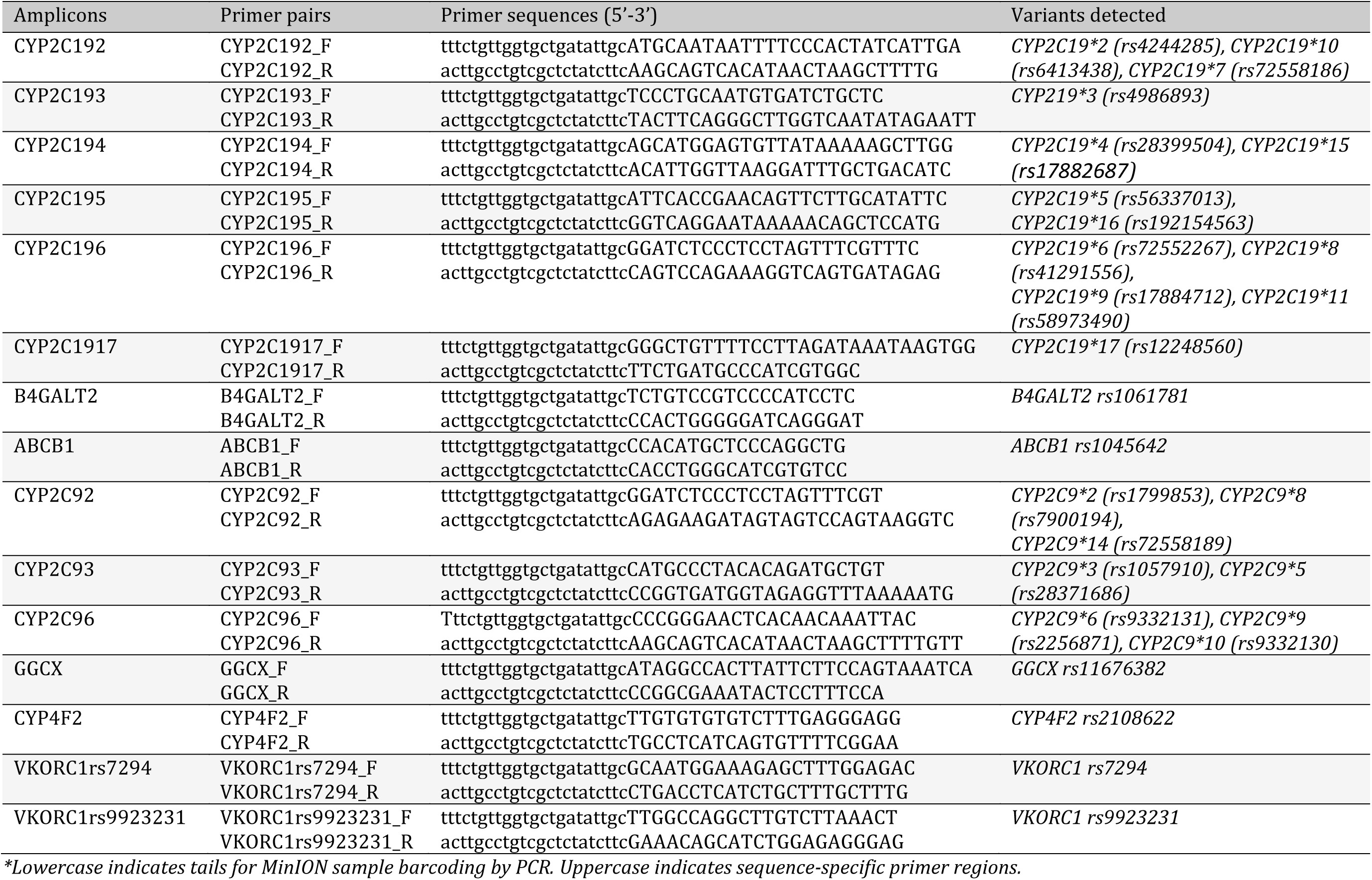
Primers and variants detected in the pharmacogenetics assay panel

## Materials and Methods

### Samples

Five reference DNA samples (Coriell Institute, Camden, NJ, USA - see Supplementary Table 1) and one in-house DNA sample with known genotypes were used as controls. Seventy eight samples from a cohort of patients who had undergone percutaneous coronary intervention (PCI) [16], with available *CYP2C19*\*2, *CYP2C19*\*3, and *CYP2C19*\*17 genotype data, were included to validate the assay.

### Multiplex PCR

A 15-plex PCR, covering a total of 27 variants, was designed using MFE-primer3.0 (https://design.igenetech.com/) (Table 1). Amplicon lengths varied between 163 – 240 bps. The forward and reverse primers were prepared separately to make 10 μM of each pool. The multiplex PCR reaction consisted of 1 μM each of the forward and reverse primer pools with 1x PCR reaction buffer, 2.5 mM of MgCl_2_, 0.4 mM of dNTPs, 1 M of betaine and 2.5 units of Taq DNA polymerase in a 50 μL reaction. The PCR was run for 35 cycles of 94 ^°^C for 15 seconds, 59 ^°^C for 15 seconds and 72 ^°^C for 30 seconds followed by a final 72 ^°^C for 1 minute. PCR product from each sample was purified using HighPrep™ PCR magnetic beads (Magbio, Gaithersburg, USA) and quantified using a Qubit dsDNA HS Assay Kit (Thermo Fisher Scientific).

### Barcoding PCR

One nM of the purified product of each sample was assigned to a specific barcode, using the PCR Barcoding Expansion 1-96 kit (EXP-PBC096) (ONT) by a second-round PCR, according to the 1D PCR Barcoding (96) amplicon (SQK-LSK108) protocol (ONT). Reactions consisted of 1x PCR buffer, 1.5 mM of MgCl_2_, 0.2 mM of dNTPs, 0.2 μM of each barcode, and 5 units of Taq DNA polymerase in 100 μL PCR reaction. The PCR consisted of 20 cycles of 94 ^°^C for 15 seconds, 62 ^°^C for 15 seconds, and 72 ^°^C for 30 seconds with final extension at 72 ^°^C for 1 minute. Taq Ti DNA polymerase (Fisher Biotec, Wembley, Australia) was used for both the initial multiplex PCR and the barcoding PCR.

### MinION library preparation

The barcoded samples were purified using HighPrep™ PCR magnetic beads (Magbio, Gaithersburg, USA). Each barcoded sample was quantified with Qubit, normalized in concentration and pooled in equimolar amounts. Eighty ng of the pooled DNA was used in the MinION library preparation, according to the 1D PCR Barcoding (96) amplicons (SQK-LSK108) protocol (ONT).

### Data analysis

The MinION sequencing was run for 48 hours as per protocol using an R9 flowcell. The fast5 files generated were basecalled and demultiplexed using Albacore version v2.3.0 (ONT), resulting in a fastq file for each barcode. The fastq files were aligned to a custom reference sequence which was built from human hg19 reference sequences of each amplicon in a fasta file (Supplementary file) using BWA MEM. Three different variant calling tools, marginCaller from marginAlign [17], nanopolish [18], and VarScan 2 [19] were used to call variants. NanoOK [20], Minion-QC [21] and Qualimap [22] were used to generate quality control parameters including read errors, read depth, and percentage of reads alignment. Integrative Genomics Viewer (IGV, Broad Institute, Boston, MA)[23] was used to visualise alignment files. Read depth plot was generated using ggplot2 in R (Vienna, Austria).

### Sanger sequencing

A nested PCR using 0.5 μM of sequence-specific primers was performed in a 20 μL reaction containing 1x PCR buffer, 1.5 mM MgCl_2_, 0.2 mM dNTPs, 0.5 units of Taq polymerase and 1 μL of 10x diluted multiplex PCR product. PCR consisted of 20 cycles of 94 ^°^C for 15 seconds, 59 ^°^C for 15 seconds, and 72 ^°^C for 30 seconds with final extension at 72 ^°^C for 1 minute. Sanger sequencing was performed using 1 μL product of nested PCR, 1 μL of primer, 1x sequencing buffer, and 0.5 μl of Big Dye Terminator in 10 μL reaction. Dye incorporation was performed in a thermal cycler with 25 cycles of 96 ^°^C for 10 seconds, 50 ^°^C for 5 seconds, and 60 ^°^C for 4 minutes, and subsequently purified using Sephadex G-50 before being sequenced using the AB3130xL genetic analyzer (Thermo Fisher Scientific).

## Results

### Read characteristics

One sequencing run containing 84 barcoded samples produced 0.9 million reads, of which one third passed the Albacore default quality threshold qscore of seven. These reads were then used for further downstream analysis. The majority of the passed reads (94.6 %) mapped to the reference sequence. The average error rate was 2.94%, 3.83%, and 3.82% for insertion, deletion, and substitution errors, respectively.

The average read depth of each amplicon varied significantly between 43 and 985x (F(14,1200)=146.6, p<0.001). Variability was also significantly different among amplicons (Levene’s test P < 0.001) (Figure 1). Three amplicons, ABCB1, CYP2C96 and VKORC1rs7294 showed very high read depth, while four amplicons, CYP2C192, CYP2C1917, CYP2C194, and CYP2C93 showed relatively low read depth of < 100 across all samples. Three out of the 78 clinical samples were excluded from analysis due to low yield in initial PCR and low read depth across all amplicons, with less than 50 % of reads being mapped to the reference sequence. Twelve samples for B4GALT2, two samples for CYP2C194, one sample for CYP2C93, and four samples for CYP4F2 amplicons had very low read depth (<10), and were also excluded from analysis (Supplementary Table 1).

**Figure 1.**
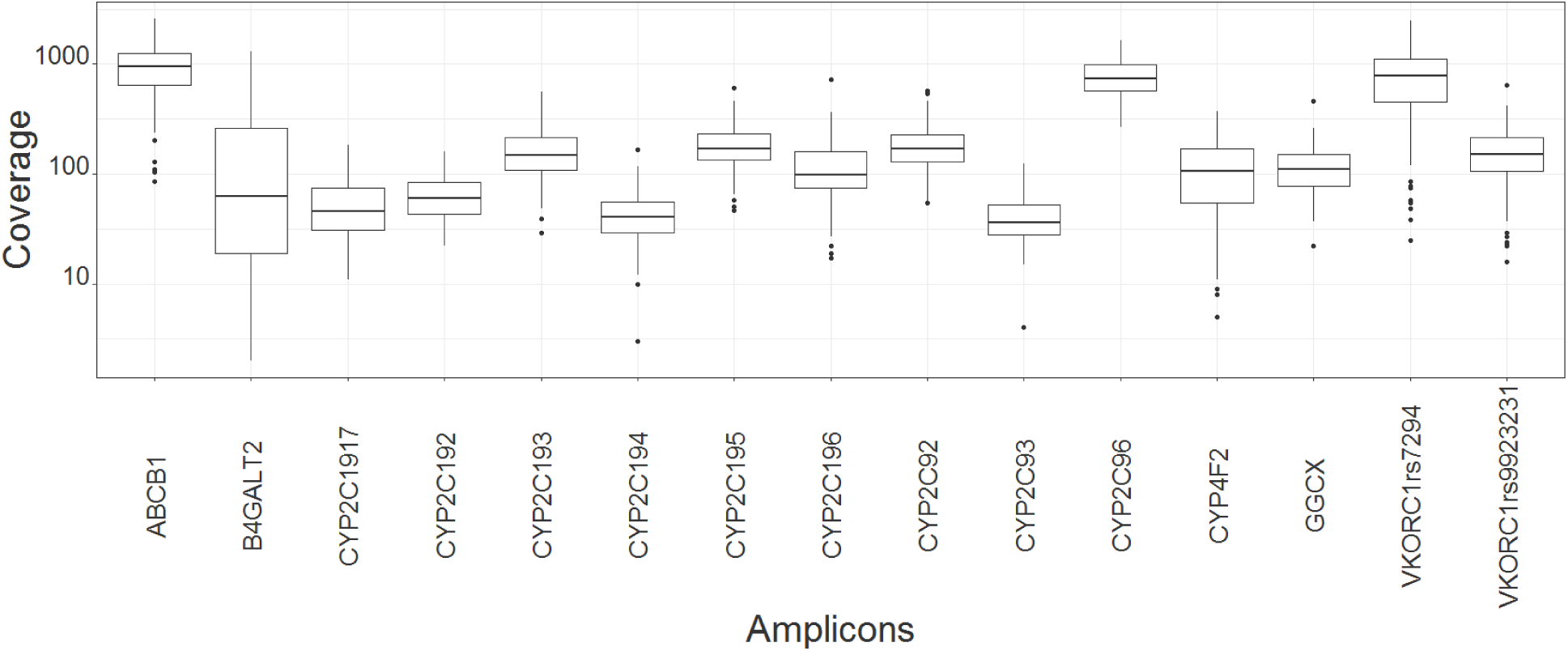
Boxplot of read depth of amplicons

### Genotyping accuracy across different variant calling tools

#### *CYP2C19*\*2, *CYP2C19*\*3, and *CYP2C19*\*17

For three genotypes (*CYP2C19*\*2, *CYP2C19*\*3, and *CYP2C19*\*17), data from two orthogonal assay platforms (Nanosphere Verigene^®^ or Sequenom MassARRAY platform) were available [16], and initial comparative analysis was done on these three genotypes. Accuracy for *CYP2C19*\*3 genotyping was 100 % across all variant calling tools (Figure 2), with the three heterozygote samples for *CYP2C19*\*3 being accurately detected using the MinION assay.

**Figure 2.**
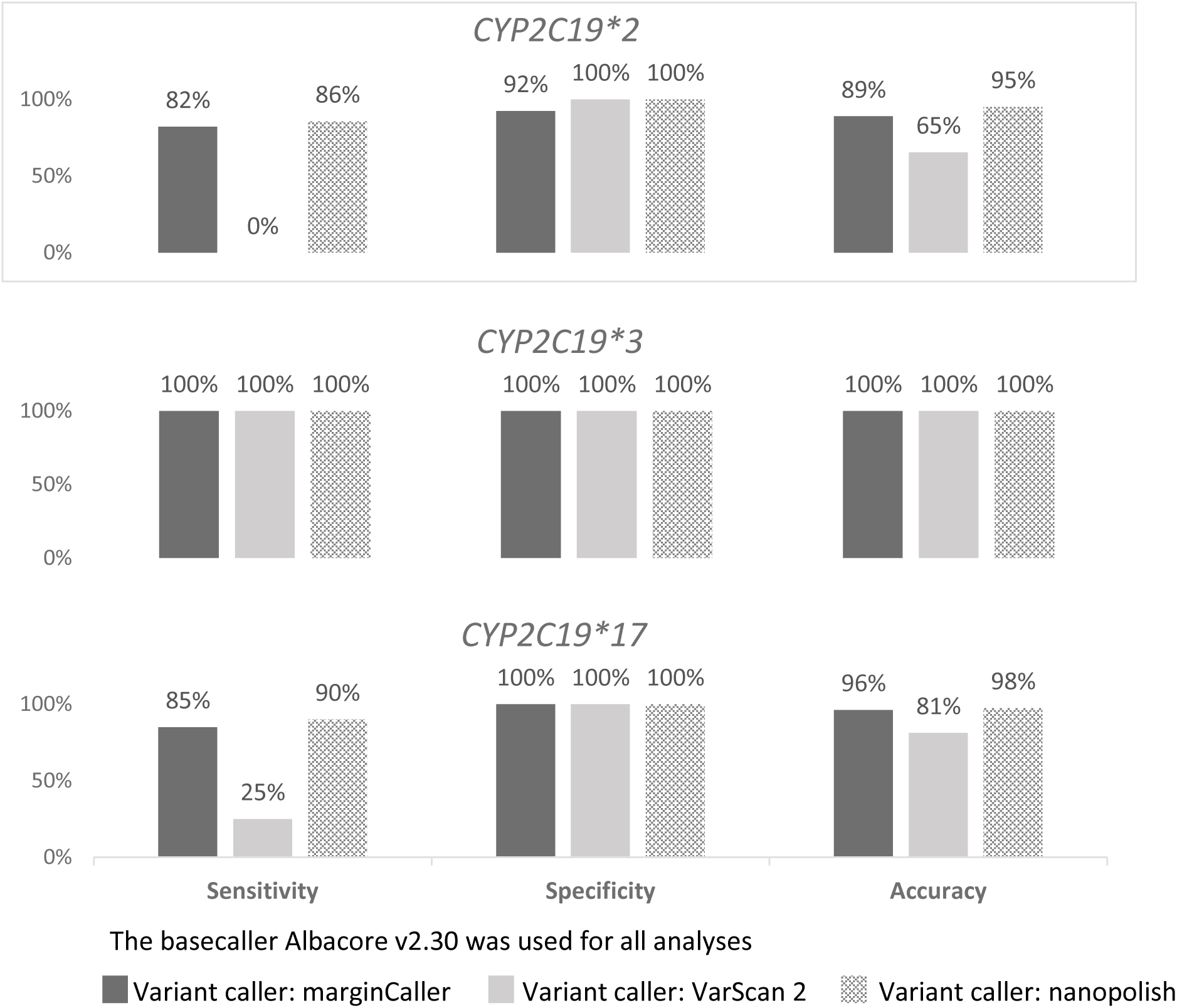
Genotyping accuracy across different bioinformatics workflow for three CYP2C19 SNPs. Threshold for heterozygote variants was set at 0.2 and homozygote at 0.65, except for nanopolish, where there is no threshold for homozygote. Nanopolish uses its own algorithm to call a variant as homozygote based on the score of the variant compared to its reference sequence (https://github.com/jts/nanopolish). Homozygote variants mis-genotyped as heterozygote were considered false negative.

Accuracy of *CYP2C19*\*2 and *CYP2C19*\*17 calls with the MinION varied depending on the bioinformatic tools used to base-call and generate the variant genotype calls. Nanopolish was the most accurate variant calling tool for this assay, with only four discordant genotypes for *CYP2C19*\*2 and two for *CYP2C19*\*17 across a total of 81 samples. All four discordant samples for *CYP2C19*\*2 and one discordant sample for *CYP2C19*17* were of low read depth (≤ 20). The other discordant sample for *CYP2C19*\*17 resulted from inaccurate calling of a homozygote variant as a heterozygote. The fraction of total reads that support the alternate allele for this sample was close to 1 (0.78), indicating a homozygote variant. Moreover, the read depth was > 100, however nanopolish incorrectly called this sample as heterozygote.

### Other variants

Analysis of the other variants included in this assay was first done on control samples. Nanopolish resulted in 100% accuracy on control samples while marginCaller missed the CYP2C9*6 variant in one sample and VarScan 2 missed a total of 11 variants in six control samples (Supplementary Table 1). As nanopolish gave the best accuracy compared to the other two variant calling tools, it was used to analyze other variants across all samples. Many of the variants included in our custom designed assay are rare, and there were only eight variants which were detected in at least three of the 81 samples. We performed Sanger sequencing for the eight variants for randomly selected subsets of the samples. Accuracy of genotype data from MinION sequencing compared with Sanger sequencing is shown in Table 2.

**Table 2.**
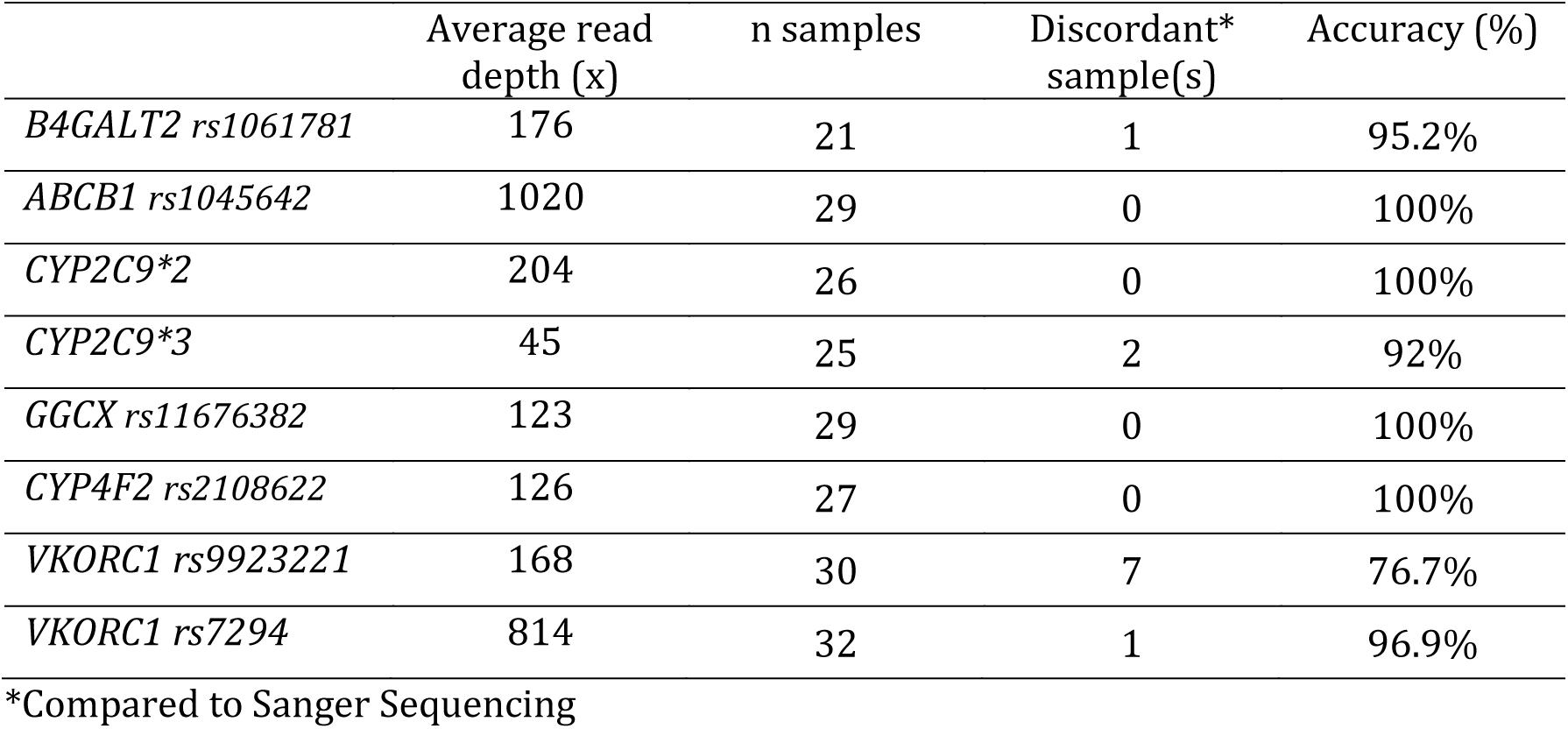
Genotyping accuracy of eight variants found in the samples

|  | Average read<br>depth (x) | n samples | Discordant*<br>sample(s) | Accuracy (%) |
| --- | --- | --- | --- | --- |
| *B4GALT2 rs1061781* | 176 | 21 | 1 | 95.2% |
| *ABCB1 rs1045642* | 1020 | 29 | 0 | 100% |
| *CYP2C9\*2* | 204 | 26 | 0 | 100% |
| *CYP2C9\*3* | 45 | 25 | 2 | 92% |
| *GGCX rs11676382* | 123 | 29 | 0 | 100% |
| *CYP4F2 rs2108622* | 126 | 27 | 0 | 100% |
| *VKORC1 rs9923221* | 168 | 30 | 7 | 76.7% |
| *VKORC1 rs7294* | 814 | 32 | 1 | 96.9% |

\*Compared to Sanger Sequencing

Two heterozygote samples for *CYP2C9*\*3 were called as wild-type by nanopolish. These two samples were of relatively low read depth (< 30). The discordant samples for *B4GALT2*, *VKORC1* rs9923231, and *VKORC1* rs7294 variants consisted of homozygote samples which were mis-called as heterozygotes. This inaccuracy could not be explained by read depth, as some of these samples had a read depth > 100. Genotype data for all samples is available in Supplementary Table 1.

The discordant results for *VKORC1* rs9923231 could be resolved by analyzing the BAM files. As seen in Figure 3, mis-called homozygote samples (lines 1 and 2) had distinct alternate allele frequency compared to real heterozygote samples (lines 3 and 4) and could be easily differentiated.

**Figure 3.**
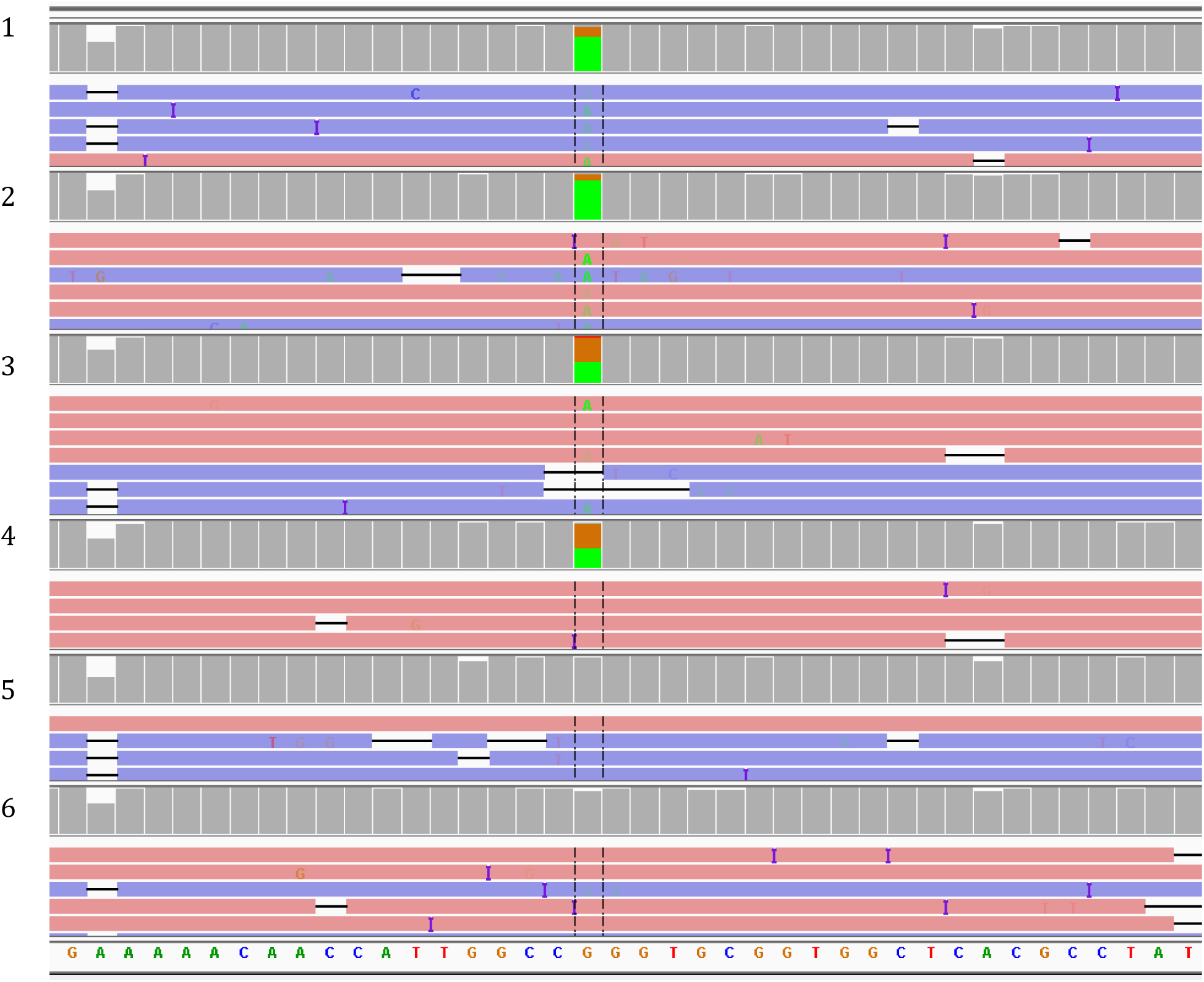
Bam files of homozygote (AA) samples (line 1 and 2), heterozygote (GA) samples (line 3 and 4) and wild-type (GG) samples (line 5 and 6) of VKORC1 rs9923231, as visualized in IGV.

### *CYP2C9*\*6, a single base deletion

All the variants included in the assay are single base substitutions, except for *CYP2C9*\*6 which is a single base deletion (delA). This variant is very rare in populations other than African, where the allele frequency is approximately 1% [24]. We included a reference sample containing a heterozygote *CYP2C9*\*6 variant. Nanopolish and VarScan 2 successfully detected this variant, while marginCaller failed. This showed that when using an appropriate variant calling tool, a single base deletion can be accurately detected in MinION sequencing data.

## Discussion

Drug responses are often affected by multiple variants in multiple genes, and detection of such variants using conventional approaches like Sanger sequencing or single PCR is both laborious and time-consuming. Extensive genotyping panels containing high number of variants can be analysed in advanced instruments like next generation sequencers, or multiplex PCR-based technologies with bead-based or mass spectrometry detection. However, these approaches require high-capital instrument costs which are not always accessible by diagnostic laboratories. Moreover, modification of such genotyping panels might be difficult and expensive, as special primers or probes are needed. The MinION on the other hand is a portable sequencing tool, available to any laboratory scale and has been use in field settings as far as outer space [18, 25, 26].

We have described an assay detecting multiple genetic variants known to affect response to clopidogrel and/or warfarin therapy. This assay can easily be modified by designing new sets of primers. The capability of multiplexing up to 96 samples in one sequencing run resulted in a reasonably cost-effective approach to genotyping. Cost-effectiveness might further be improved with the smaller-scale, cheaper “Flongle” flow cells recently released by ONT.

The simple workflow of only one PCR reaction per sample was achieved by designing short amplicons, which were previously shown to be compatible with the MinION [27]. All contigs were amplified successfully, and despite the uneven yield among amplicons, the majority of genotypes could be accurately determined using appropriate bioinformatics tools. Genotyping accuracy was in part dependent on the variant calling tool. We showed that nanopolish achieved the best accuracy compared with marginCaller and VarScan 2. Instead of calling variants as single bases, nanopolish investigates a block of 10 bases at a time, and calculates the best likely haplotype using a hidden Markov model [18]. Nanopolish could detect both single base substitutions and a single base deletion.

Although generally accurate, a small proportion of incorrect genotype calls were observed. The majority of these were homozygote variants miscalled as heterozygotes. As previously reported, this seems to be a MinION and/or nanopolish specific problem [10]. Using nanopolish to genotype a fraction of variants on chromosome 20, the authors reported that 3,217 out of 4,781 mislabelled variants were due to this kind of error [10]. Improvements to variant calling tools for nanopore sequencing data may lead to more accurate calling.

Accuracy in MinION sequencing is also sequence-dependent [28]. This was well reflected in this assay with some genotypes showing better accuracy than the others. In particular, *CYP2C19*\*3 showed perfect accuracy regardless of the bioinformatic tools used to genotype. *VKORC1* rs9923231, on the contrary, showed the least accurate genotyping despite the relatively high read depth (> 100). However, these errors could be resolved by visual inspection of the alignment (BAM) file.

## Conclusion

Drug-specific or broad pharmacogenetic assays are possible on the MinION sequencer, with high numbers of amplicons and sample multiplexing. High accuracy was achieved for most of the variants in our custom genotyping panel by applying suitable bioinformatic tools. The choice of variant calling tool is important, with nanopolish performing best for the assay described here. However, some mis-genotyping issues remain, which need to be resolved before such assays could be applied in a clinical or diagnostic setting. Continual improvements in chemistry and flowcells by ONT, and continuing development of bioinformatic tools by the research community is likely to resolve these issues in the future.

