## Supplementary file for "A Multiplex Pharmacogenetics Assay using the MinION Nanopore Sequencing Device"

Reference Sequence

>CYP2C1927

ATGCAATAATTTTCCCACTATCATTGATTATTTCCCGGGAACCCATAACAAATTACTTAAAAACCTTGCTTTTATGGAAAGTGATATTTTGGAGAAAGTAAAAGAACACCAAGAATCGATGGACATCAACAACCCTCGGGACTTTATTGATTGCTTCCTGATCAAAATGGAGAAGGTAAAATGTTAACAAAAGCTTAGTTATGTGACTGCTT

>CYP2C193

TCCCTGCAATGTGATCTGCTCCATTATTTTCCAGAAACGTTTCGATTATAAAGATCAGCAATTTCTTAACTTGATGGAAAAATTGAATGAAAACATCAGGATTGTAAGCACCCCCTGGATCCAGGTAAGGCCAAGTTTTTTGCTTCCTGAGAAACCACTTACAGTCTTTTTTTCTGGGAAATCCAAAATTCTATATTGACCAAGCCCTGAAGTA

>CYP2C194

AGCATGGAGTGTTATAAAAAGCTTGGAGTGCAAGCTCACGGTTGTCTTAACAAGAGGAGAAGGCTTCAATGGATCCTTTTGTGGTCCTTGTGCTCTGTCTCTCATGTTTGCTTCTCCTTTCAATCTGGAGACAGAGCTCTGGGAGAGGAAAACTCCCTCCTGGCCCCACTCCTCTCCCAGTGATTGGAAATATCCTACAGATAGATATTAAGGATGTCAGCAAATCCTTAACCAATGT

>CYP2C195

ATTCACCGAACAGTTCTTGCATATTCTGTCTGTGCCAGTTATAGAGACAGTGTTTGTCACTCTCACAGTTACACATGAGGAGTAACTTCTCCCTATGTTTGTTATTTTCAGGAAAACGGATTTGTGTGGGAGAGGGCCTGGCCCGCATGGAGCTGTTTTTATTCCTGACC

>CYP2C1968

GGATCTCCCTCCTAGTTTCGTTTCTCTTCCTGTTAGGAATCGTTTTCAGCAATGGAAAGAGATGGAAGGAGATCCGGCGTTTCTCCCTCATGACGCTGCGGAATTTTGGGATGGGGAAGAGGAGCATTGAGGACCGTGTTCAAGAGGAAGCCCGCTGCCTTGTGGAGGAGTTGAGAAAAACCAAGGGTGGGTGAACATACTCTCTATCACTGACCTTTCTGGACTG

>CYP2C1917

GGGCTGTTTTCCTTAGATAAATAAGTGGTTCTATTTAATGTGAAGCCTGTTTTATGAACAGGATGAATGTGGTATATATTCAGAATAACTAATGTTTGGAAGTTGTTTTGTTTTGCTAAAACAAAGTTTTAGCAAACGATTTTTTTTTTCAAATTTGTGTCTTCTGTTCTCAAAGCATCTCTGATGTAAGAGATAATGCGCCACGATGGGCATCAGAA

>B4GALT2

TCTGTCCGTCCCCATCCTCAGGATCTCCCTGACTGGGATGAAGATCTCACGCCCAGACATCCGAATCGGCCGCTACCGCATGATCAAGCACGACCGCGACAAGCATAACGAACCTAACCCTCAGAGGTGACCCCAGCACCCTCACCCCTTACTCCCCAGAGGCAACTTCCCAATATCCCCAACTCTTGACCCCAAGTGGCCCAATCCCTGATCCCCCAGTGG

>ABCB1

CACCTGGGCATCGTGTCCCAGGAGCCCATCCTGTTTGACTGCAGCATTGCTGAGAACATTGCCTATGGAGACAACAGCCGGGTGGTGTCACAGGAAGAGATTGTGAGGGCAGCAAAGGAGGCCAACATACATGCCTTCATCGAGTCACTGCCTAATGTAAGTCTCTCTTCAAATAAACAGCCTGGGAGCATGTGG

>CYP2C92

GGATCTCCCTCCTAGTTTCGTTTCTCTTCCTGTTAGGAATTGTTTTCAGCAATGGAAAGAAATGGAAGGAGATCCGGCGTTTCTCCCTCATGACGCTGCGGAATTTTGGGATGGGGAAGAGGAGCATTGAGGACCGTGTTCAAGAGGAAGCCCGCTGCCTTGTGGAGGAGTTGAGAAAAACCAAGGGTGGGTGACCCTACTCCATATCACTGACCTTACTGGACTACTATCTTCTCT

>CYP2C93

TGCCCTACACAGATGCTGTGGTGCACGAGGTCCAGAGATACATTGACCTTCTCCCCACCAGCCTGCCCCATGCAGTGACCTGTGACATTAAATTCAGAAACTATCTCATTCCCAAGGTAAGTTTGTTTCTCCTACACTGCAACTCCATGTTTTCGAAGTCCCCAAATTCATAGTATCATTTTTAAACCTCTACCATCACCGG

>CYP2C96

CCCGGGAACTCACAACAAATTACTTAAAAACGTTGCTTTTATGAAAAGTTATATTTTGGAAAAAGTAAAAGAACACCAAGAATCAATGGACATGAACAACCCTCAGGACTTTATTGATTGCTTCCTGATGAAAATGGAGAAGGTAAAATGTAAACAAAAGCTTAGTTATGTGACTGCTT

>CYP4F2

TGCCTCATCAGTGTTTTCGGAACCCATCACAACCCAGCTGTGTGGCCGGACCCTGAGGTGCGGGGCCCCTCTCTCTGTTTTTGTCCATTCCAAGGCTCCTAGAGGAGGGGGCAGGGTTTTGATCAGGAGTATCCAACATCACCTCCCTCAAAGACACACACAA

>GGCX

CCGGCGAAATACTCCTTTCCATGAGCGATTCTTCCGCTTCTTGTTGCGAAAGCTCTATGTCTTTCGCCGCAGGTAAGTTCACAACAATATTTGTCATTGCCATCATATGTTGGCAAGCTTGGTAACTTTCCCCTGGGGAGAGTAACTCTAGAGACTGTCATACAGGAAAGGACAGTTTAGCTCTAGGTATCTTTTCCTGCCATTCTTCTTGATTTACTGGAAGAATAAGTGGCCTAT

>VKORC1rs7294

CTGACCTCATCTGCTTTGCTTTGGCATGTGAGCCTTGCCTAAGGGGGCATATCTGGGTCCCTAGAAGGCCCTAGATGTGGGGCTTCTAGATTACCCCCTCCTCCTGCCATACCCGCACATGACAATGGACCAAATGTGCCACACGCTCGCTCTTTTTTACACCCAGTGCCTCTGACTCTGTCCCCATGGGCTGGTCTCCAAAGCTCTTTCCATTGC

>VKORC1rs9923231

GAAACAGCATCTGGAGAGGGAGGAGCCAGCAGGAGAGGGAAATATCACAGACGCCAGAGGAAGAGAGTTCCCAGAAGGGTAGGTGCAACAGTAAGGGATCCCTCTGGGAAGTCAAGCAAGAGAAGACCTGAAAAACAACCATTGGCCGGGTGCGGTGGCTCACGCCTATAATCCTAGCATTTTGGGAGGCCGAGGTGGGTGGATCACTTGAGGTCAGGAGTTTAAGACAAGCCTGGCCAA
